# Long-range Diffusion of Bound Ca^2+^: Time-dependent WKBJ solutions and in One- and Two-Dimensional Imaging

**DOI:** 10.64898/2026.07.06.735233

**Authors:** S. L. Mironov

**Affiliations:** Institute of Neuro- and Sensory Physiology, Georg-August-University, Göttingen 37073, Germany

**Keywords:** Cytoplasmic calcium profiles, buffer motility, synuclein channels, L-type Ca^2+^ channels, propagating calcium signals, analytical solutions, WKBJ

## Abstract

Reaction–diffusion (RD) systems play a fundamental role in numerous biochemical and biophysical processes. Here, we present a novel analytical framework for solving RD equations by applying the Wentzel–Kramers–Brillouin–Jeffreys (WKBJ) formalism to Ca^2+^ buffering. Previous models have primarily focused on stationary Ca^2+^ nanodomains while neglecting diffusion and saturation of intracellular Ca^2+^ buffers and sensors. We derive analytical solutions using WKBJ ansatz and demonstrate that sustained Ca^2+^ source produces continuously expanding distributions of free Ca^2+^, whereas the concentration of Ca^2+^-bound buffers develops at much slower timescale. These predictions are supported experimentally by measurements of one-dimensional fluorescence profiles produced by single-channel activity and two-dimensional profiles generated by whole-cell Ca^2+^ currents. The analytical framework developed here extends Michaelis–Menten-type kinetics to reaction–diffusion systems and may therefore be broadly applicable to biochemical and biophysical processes in which diffusion cannot be neglected.

## 1 Introduction

The Michaelis–Menten equation (1913) and its associated quasi-steady-state approximation (Segel and Slemrod, 1989; Berzani et al. 2015; Choi et el. 2017) have served for more than a century as fundamental tools for describing biochemical and biophysical reactions. Despite numerous refinements, these formulations generally describe either spatially homogeneous systems or highly localized reactions in which diffusion is neglected.

For intracellular Ca^2+^ signaling, Wagner and Keizer (1994) incorporated Michaelis–Menten kinetics (called also rapid buffer approximation, RBA, for Ca^2+^ buffering) into a reaction–diffusion framework and derived a nonlinear equation for total Ca^2+^ concentration, which they analyzed numerically and under several limiting approximations. Mironova & Mironov (2008) found that total Ca^2+^ can be well described using the logarithmic approximation and derived analytical formulae for transient free Ca^2+^ dynamics. They also emphasized that Ca^2+^ is transported predominantly in its bound form within the cytoplasm.

Many endogenous Ca^2+^ buffers diffuse slowly or are effectively immobilized (Zhou & Neher, 1993; Schwaller, 2010). Mironov (2023a) analyzed the case for irreversible Ca^2+^ binding and derived analytical expressions that predicted a localized accumulation of free Ca^2+^ around single channel.

Ca^2+^-bound buffer species were found to propagate away from it much faster. Because endogenous buffers and sensors are usually large proteins with limited mobility, the propagation of bound Ca^2+^ may represent an efficient mechanism for long-range intracellular signaling. In this view, Ca^2+^-bound molecules, rather than free Ca^2+^ itself, may constitute the principal carriers of information between distant intracellular sites.

At first sight, the hypothesis may be thought to be inconsistent with the widely accepted concept of stationary Ca^2+^ nanodomains surrounding single channels (Neher, 1986; Eisner et al. 2023).

However, these perspectives are not mutually exclusive, as discussed below.

In the present study, we revisit Michaelis–Menten-type kinetics by explicitly incorporating diffusion and derive analytical solutions using the Liouville–Green transformation, commonly known in quantum mechanics as the Wentzel–Kramers–Brillouin–Jeffreys (WKBJ) approximation. Although WKBJ methods are routinely applied to stationary problems, their use in non-stationary reaction– diffusion systems has remained limited (Krause et al. 2011).

To maintain the accessibility of the main text, all mathematical derivations are presented in the Sections 5.1 and 5.2. We demonstrate that the leading-order term of the Liouville–Green/WKBJ expansion accurately reproduces multidimensional diffusion with and without reactions and provides simple analytical expressions for Ca^2+^ buffering. Experimental measurements obtained in one-dimensional single-channel recordings and two-dimensional neuronal preparations support the theoretical predictions.

Overall, our findings suggest that Ca^2+^-bound species can propagate more rapidly and over greater distances than localized free Ca^2+^ signals, potentially influencing the timing and spatial organization of intracellular signaling. Beyond Ca^2+^ dynamics, the analytical framework developed here offers a straightforward approach for incorporating diffusion into Michaelis–Menten-type reaction systems.

## 2. Methods

### 2.1 Ethical approval

All animal experiments were approved by the Lower Saxony State Office for Consumer Protection and Food Safety and conducted in accordance with institutional and national guidelines.

### 2.2 Cell preparation

Cultured hippocampal neurons and chick dorsal root ganglion (DRG) neurons were prepared as previously described (Mironov, 1995; Mironov et al. 1993, respectively). DRG neurons were used 6–12 h after plating, when they had fully adhered to the substrate and exhibited a flattened circular morphology with a thickness of approximately 2–3 μm. At this time in culture, spontaneous synaptic activity was absent, minimizing contributions from neuronal network activity.

### 2.3 Solutions

In single-channel measurements the solutions contained 30 mM Tris buffer (pH 7.4) together with either 154 mM NaCl (pipette) or 88 mM CaCl_2_ (bath). Fluo-4 and Fluo-4 dextran (30 kDa) were used at final concentrations of 0.2 mM. In whole-cell recordings intrapipette solution contained CsMeSO_4_ (92 mM), CsCl (43 mM) TEA-Cl (5 mM), MgCl_2_ (1 mM), HEPES (10 mM), ATP (4 mM), GTP (0.4 mM); pH 7.4 (titrated with CsOH) was used. The indicators were added at 200 µM.

### 2.4 Patch-clamp recording and imaging

Electrophysiological recordings and fluorescence imaging were performed as previously described. α-Synuclein channels were incorporated into inside-out membrane patches excised from cultured hippocampal neurons (Mironov, 2015). Opening of individual channels generated outward Ca^2+^ currents that produced localized increases in Ca^2+^-bound fluorescence within the recording pipette.

In whole-cell recordings, voltage steps were applied from −60 to −10 mV and evoked Ca^2+^ currents predominantly carried by L-type Ca^2+^ channels.

Fluorescence imaging was performed using total internal reflection fluorescence microscopy (TIRF) combined with high-speed EMCCD acquisition. In whole-cell recordings experimental protocol - control - BayK 8044 - Nifedipine (see Methods) - was applied, when L-type current showed no significant run-down within first 5 minutes. For imaging, an upright microscope (Axioscope 2, Zeiss) with a 63x objective lens (N. A. 1.4) was used. The laser beam delivered 488 nm light from below through a prism (TIRF Labs, Inc. Cary NC) at an angle appropriate to evoke TIRF. The emission (525 nm) passed through dichroic mirror filtered at 535 nm. The images are captured by a cooled CCD camera iXON Ultra 888 EMCCD (ANDOR, Oxford Instruments) operated under ANDOR software. In 1D-experiments the line scan mode (512 x 2 pixels at 12-bit resolution) was used that corresponds to 0.1 ms-time acquisition in Crop mode. In whole-cell recordings, the recorded area was 128 x 128 pixels and acquisition time was 1.5 ms. The fluorescence of fluo-4 increases >30-fold after Ca^2+^ binding that minimized out-of-focus effects. The lateral resolution measured with 0.1 um TetraSpeck beads (ThermoFisher, Germar, Germany) gave lateral point spread function 0.18 um TIRF depth was around 200 nm that allow to consider already flattened DRG cells in culture as plane objects and apply the results obtained for Ca^2+^ diffusion using circular geometry. Image analysis was performed offline using MetaMorph software. Images were background-subtracted, and median-filtered before analysis. Fluorescence changes are reported as relative changes (ΔF/F_0_).

### 2.5 Statistics

Each experiment was repeated in at least four cells obtained from different preparations. Data are presented as mean ± SD. Statistical significance was assessed using paired t-tests where appropriate, with significance accepted at *P* < 0.05.

#### 2.6 Reaction–diffusion model

Ca^2+^ diffusion in the presence of a single buffer was described by coupled reaction–diffusion equations

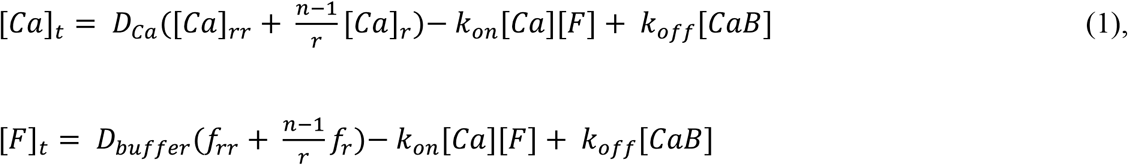

governing free Ca^2+^, free buffer (*F*), and Ca^2+^-bound (*B*) buffer concentrations. Here the Laplacian is written in *n*-dimensional form (Carslaw & Jaeger, 1986; Crank, 1979). The model incorporates diffusion coefficients and reversible binding kinetics characterized by association (*k*_*on*_) and dissociation (*k*_*off*_) rate constants representative of cytoplasmic Ca^2+^-binding proteins. The constants of Ca^2+^ buffering (Table 1) confer typical values for cytoplasmic calcium-binding proteins: *k*_*on*_ = 3^·^10^8^ M^-1^s^-1^ corresponds to diffusion-limited binding, *B*_*o*_ = 0.2 mM, and *D*_*Ca*_ = 220 μm^2^/s. Numerical values for model parameters are summarized in Table 1.

**Table 1.** Basic definitions and model parameters.

| 462 | <u>Abbreviation</u> | <u>Definition</u> | <u>Numerical value</u> |
| --- | --- | --- | --- |
| 463 | $B_o$ | Total buffer concentration | 0.2 mM |
| 464 | $c = [\text{Ca}^{2+}]/B_o$ | Normalized $\text{Ca}^{2+}$ concentration | 0.1 - 10 |
| 465 | $f = [F]/B_o$ | Normalized free buffer concentration | 0 - 1 |
| 466 | $b = [B]/B_o = 1 - f$ | Normalized $\text{Ca}^{2+}$-bound buffer concentration | 0 - 1 |
| 467 | $D_{Ca}$ | $\text{Ca}^{2+}$ diffusion coefficient | $220 \mu\text{m}^2/\text{s}$ |
| 468 | $d = D_{\text{buffer}}/D_{Ca}$ | Relative buffer diffusion coefficient | 0.03 - 1 |
| 469 | $k_{on}$ | on-rate binding constant | $3 \cdot 10^8 \text{ M}^{-1} \text{ s}^{-1}$ |
| 470 | $k_{off}$ | off-rate constant | $10^2 \text{ s}^{-1}$ |
| 471 | $K_d$ | $\text{Ca}^{2+}$-buffer dissociation constant | $0.3 \text{ } \mu\text{M}$ |
| 472 | $[\text{Ca}^{2+}]_o$ | Normalized $\text{Ca}^{2+}$ concentration at the source | 2 - 200 $\mu\text{M}$ |
| 473 | $\tau = 1/k_{on}B_o$ | Intrinsic time constant | 50 $\mu\text{s}$ |
| 474 | $r_o = \sqrt{D_{Ca}/B_o k_{on}}$ | Intrinsic space constant | 105 nm |

Analytical solutions were derived using the Liouville–Green/WKBJ formalism (Sections 5.1 and 5.2). To leading order, the resulting expressions provide compact closed-form approximations for free Ca^2+^ and bound buffer distributions in both one-dimensional and multidimensional geometries. In 1D-case

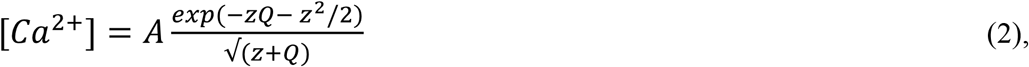

where A is the concentration of Ca^2+^ at the boundary; a self-similarity (Boltzmann) variable is *z = x/2√t, Q = √*(*z*^*2*^*+*1), and *D = √*(*1 - d*)/2 with normalized *d = D*_*buffer*_*/D*_*Ca*_. The concentration of free buffer is determined as

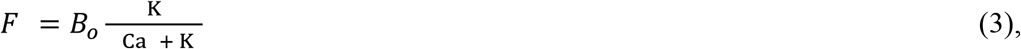

and

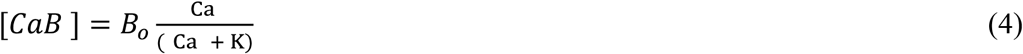

where non-dimensional dissociation constant *K = K*_*d*_*/B*_*o*_ *=* 0.003, Table 1. The expressions are in accord with RBA and MM formalism. For radial diffusion (a 3D-case), all concentrations are simply divided by *r*, and for 2D-case numerical solution for free Ca^2+^ is preferable, because analytical formulae are cumbersome. Fig. B shows that Eqs. (2 - 4) reproduce well results obtained by direct numerical solution of RD-PDE

## 3. Results

### 3.1. Theoretical prediction of one- and two-dimensional Ca^2+^ profiles

Fluorescence measurements report Ca^2+^ bound to the indicator rather than free Ca^2+^ itself and the theoretical analysis therefore focuses on the dynamics of the bound species. We used the Liouville– Green/WKBJ formalism was used to derive analytical expressions for the spatiotemporal distributions of free Ca^2+^ and Ca^2+^-bound buffer species in Section 5.2. Figure 1 compares solutions obtained for slowly diffusing buffers (relative diffusion coefficient *d* = *D*_*buffer/*_*/D*_*Ca*_ = 0.03) and more rapidly diffusing buffers (*d* = 0.8 e. g. *D*_*Fluo-4*_ = 180 µm^2^/s, Mironova and Mironov, 2008). In both cases, the leading-order WKBJ approximation closely matches the result with first-order correction added, demonstrating rapid convergence of the expansion and indicating that the leading term alone can provide a good approximation.

**Fig. 1.**
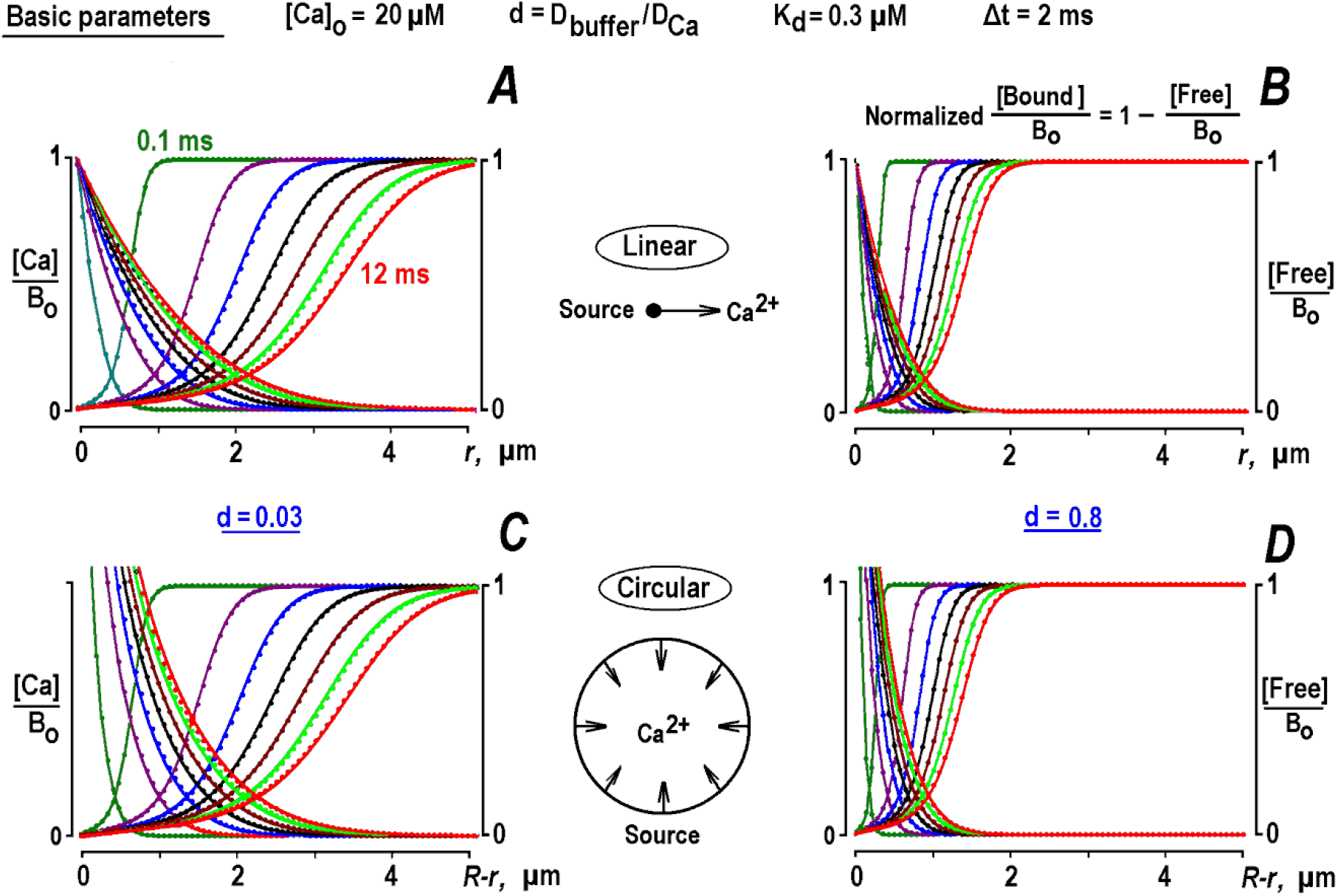
Calculated profiles of free Ca^2+^ and buffer. Time- and spatial dependencies of free Ca^2+^ and buffer. ***A, B*** - Linear diffusion from the point source. ***C, D*** - Diffusion from the surface surrounding a circular cell. Differently colored curves are placed 2 ms apart and start from 0.1 ms. They present a leading WKBJ term solution derived in Section 5.2. The dots indicate its correction in the first WKBJ order and indicate marginal contributions. Time and space scales are the same in all panels as well as in Fig. 2. In linear case the distance *r* is measured from Ca^2+^ source and in circular case the distance *R - r* is measured from the cell surface (*R* = 6 µm). The concentrations of free Ca^2+^ and buffer are normalized to the total buffer concentration, *B*_*o*_ = 0.2 mM. Left panels (***A*** and ***C***) present solutions for basic model parameters (Table 1, also enlisted in top string). In right panels (***B*** and ***D***) the relative diffusion coefficient *d* increased from 0.03 to 0.8 as indicated. Free Ca^2+^ show a steady increase that slowly depletes buffer at larger distances. Because a sum of normalized [bound Ca^2+^] and [free buffer] is equal to 1, they mirror each other and the concentration of bound Ca^2+^ can be readily obtained. Note that in both 1D- and 2D-cases the patterns of free Ca^2+^ and buffer become close at big distances from Ca^2+^ source. For circular geometry they deviate only at small distances due to effect of the term *c*_*r*_*/r* in the 2D-Laplacian and correspondingly cut off from above in the panels ***C*** and ***D***. The data are generated for a constant Ca^2+^ concentration at the source.

For slowly diffusing buffers, free Ca^2+^ spreads progressively from the source as buffer becomes locally depleted. In contrast, when buffer mobility is closer than that of free Ca^2+^, both free and bound Ca^2+^ distributions remain substantially more confined. This behavior is consistent with the narrow Ca^2+^ nanodomains typically inferred from measurements obtained with small organic fluorescent indicators, whose diffusion coefficients are only moderately lower than that of free Ca^2+^. Comparison of one-dimensional and circular geometries shows that the predicted free Ca^2+^ profiles resemble each other at distances greater than >1 μm and times exceeding 2 ms. Deviations near the source arise from the geometry-dependent Laplacian term and become negligible at larger distances.

For a sustained Ca^2+^ source, free Ca^2+^ concentrations increase continuously because progressive buffer saturation reduces local buffering capacity. In contrast, the distributions of Ca^2+^-bound and free buffer evolve as traveling fronts that propagate away from the source. Calculated velocities are similar for linear and circular geometries (Table 2), indicating that this behavior is largely independent of spatial dimensionality.

**Table 2.**
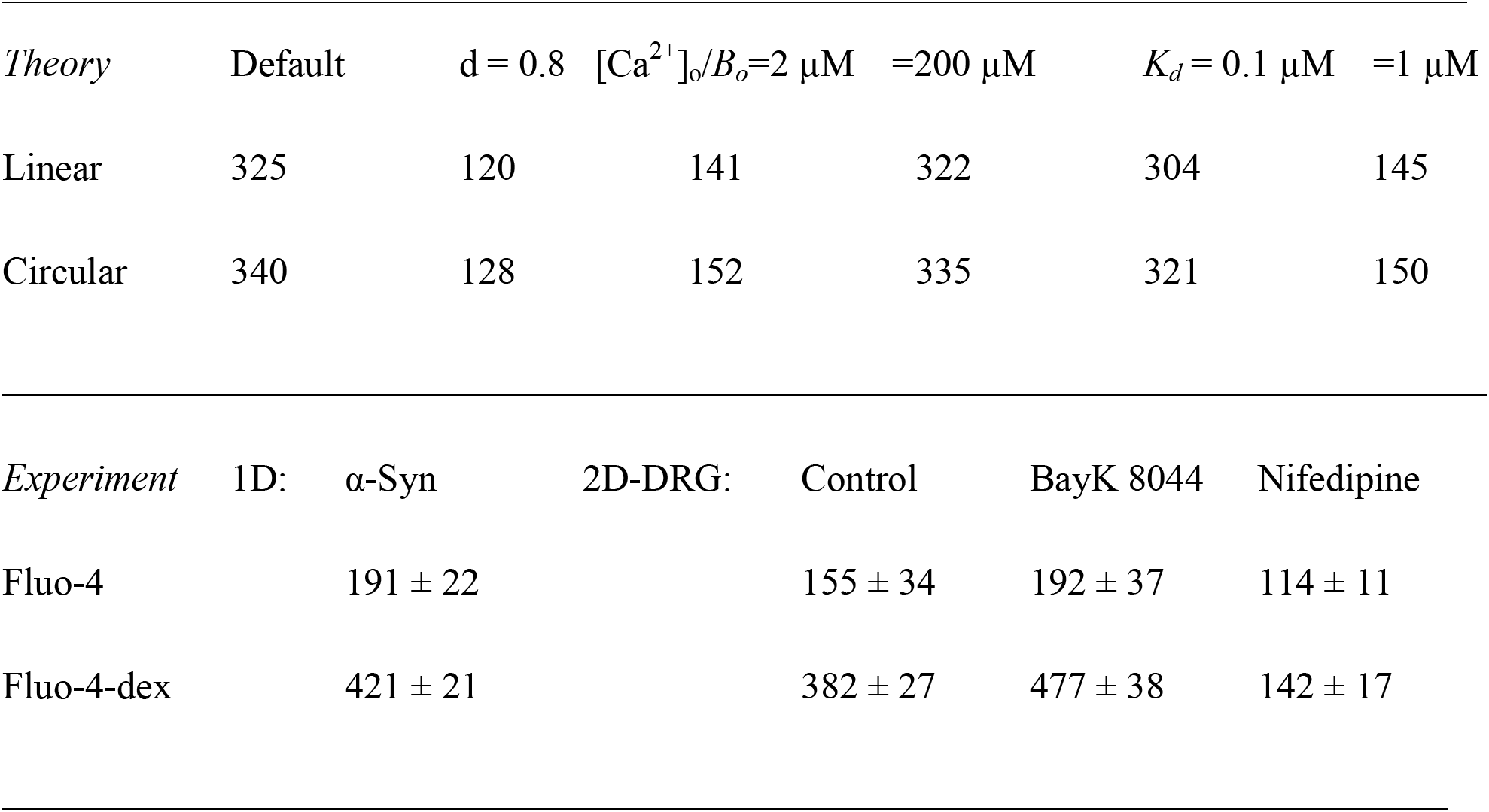
Estimation of velocities (in µm/s) of Ca^2+^ -bound buffer waves.

The calculations shown in Figure 1 assume sustained Ca^2+^ flux. For channels with shorter open times, such as voltage-gated Ca^2+^ channels, the expansion of profiles would be terminated upon channel closure and therefore acquire spatially narrower patterns. Under these conditions, the apparent width of the free Ca^2+^ domain is determined jointly by diffusion and channel open time, whereas the distribution of bound Ca^2+^ additionally reflects the kinetics of buffer binding. Consequently, the duration of Ca^2+^ influx may critically influence downstream signaling by determining whether sufficient sensor activation can occur before channel closure.

One possible test of the proposed model could be examination of correlation between the duration of opening of Ca^2+^ channels and activity of Ca^2+^-dependent channels, when present in the same patch. Previously it was noticed (Mironov, 2015) that the activity of synuclein channels did not change the activity of delayed rectifier channels (K_DR_) and modulated activity of K_ATP_ channels due to changes in submembrane ion homeostasis. It would be interesting to correlate the activity of Ca^2+^ and Ca^2+^-activated K^+^ (K_Ca_) channels. However, one must choose a preparation, where K_Ca_ channels are abundant, that is not the case for hippocampal neurons used in this study.

Figure 2 illustrates the effects of key model parameters. After normalization to the Ca^2+^ concentration at the source, the shape of the free Ca^2+^ profile is largely independent of influx amplitude. In contrast, increasing the source Ca^2+^ concentration markedly accelerates propagation of the bound species while preserving similar wave morphology in both one- and two-dimensional geometries. This behavior arises because elevated Ca^2+^ rapidly saturates local buffer molecules, causing the depletion front to propagate outward.

**Fig. 2.**
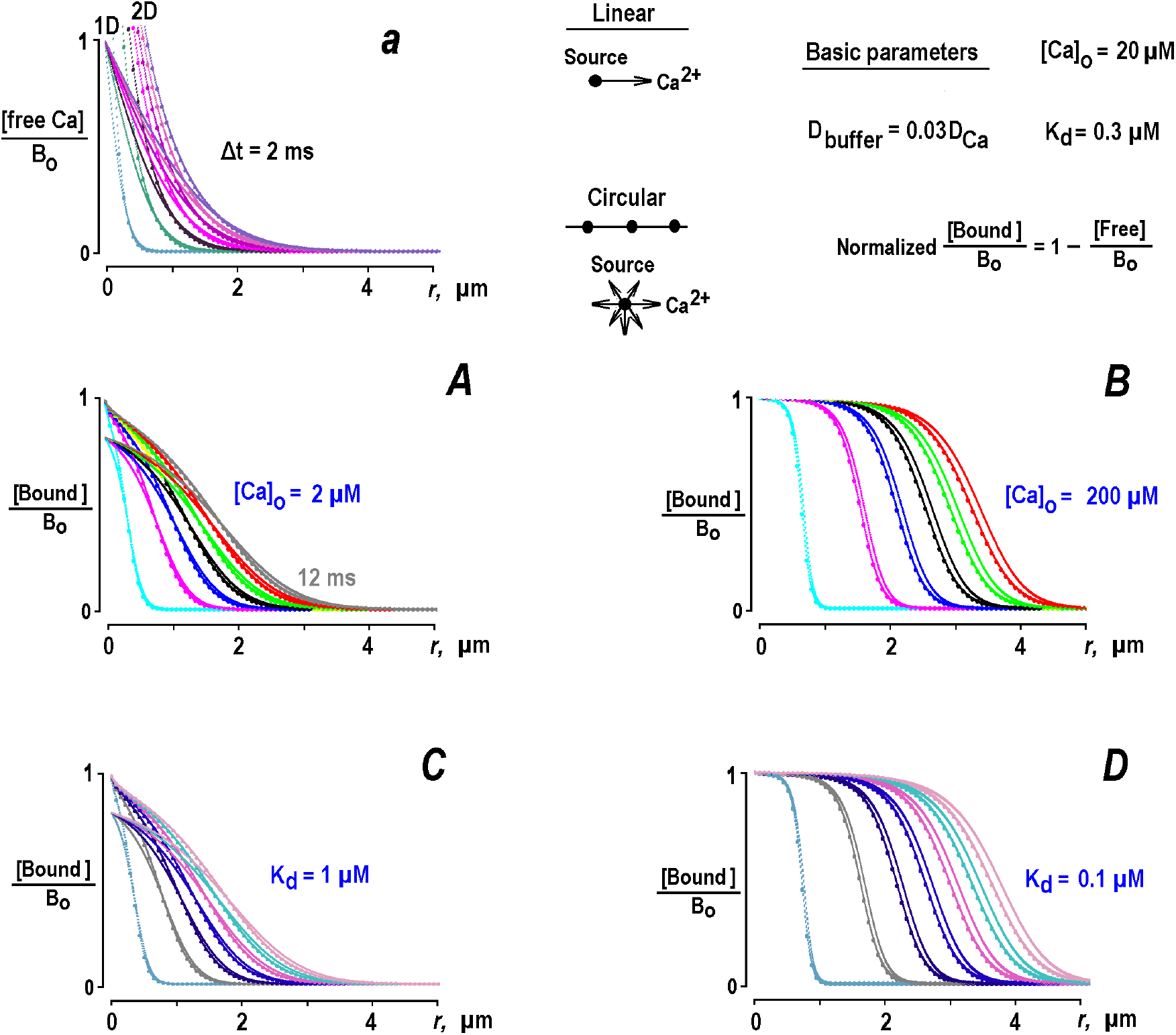
Calculated Ca^2+^ profiles for different model parameters. The data present leading term WKBJ solution derived in Section 5.2. The notations are the same as in Fig. 1. Only results for a slowly moving buffer (*d* = 0.03) are presented. ‘Default’ model parameters (Table 1), are shown also in right top corner. In each calculation, only one key parameter, [Ca^2+^]_o_ or *K*_*d*_, is modified. Individual curves start from 0.1 ms and plotted at 2 ms time interval. Free Ca^2+^ diffused from the point source (inset in the middle) and 1D- and 2D-profiles are shown only once in top left (**a**). In all cases presented they scaled proportionally according to [Ca^2+^]_o_ at the source. Initial free Ca^2+^ is much bigger in 2D-geometry. At distances >1 µm, and times >2 ms, the results for 1D-(lines) and 2D-geometries (lines with circles) practically coalesce. For other parameters, the profiles for Ca^2+^-bound buffer changed: for small [Ca^2+^]_o_ = 2 μM (***A***), bound Ca^2+^ at big distances follows expansion of free Ca^2+^(***a***), but showed slightly slower decay. At bigger [Ca^2+^]_o_= 200 μM, the pattern of bound Ca^2+^ is more extended, keep on increasing after free Ca^2+^ long subsides (***B***). For *K*_*d*_ = 0.1 μM (***D***), bound [Ca^2+^] propagates faster, whereas for bigger dissociation constant (*K*_*d*_ = 1 μM, ***C***), the profiles of bound [Ca^2+^] are more dense, resembling those obtained with small [Ca^2+^]_o_ = 2 μM (***B***).

The dissociation constant likewise influences wave propagation. Lower *K*_*d*_ values produce faster-moving fronts, whereas larger *K*_*d*_ values yield slower and more spatially confined propagation owing to more rapid equilibration between free and bound states. Across all parameter combinations examined, however, the qualitative propagating behavior of Ca^2+^-bound species remains robust.

The sigmoidal shape of the propagating fronts follows directly from the analytical solution. Under the rapid-buffer approximation, the concentration of bound buffer can be expressed as a Michaelis– Menten-type function of free Ca^2+^, Eq. (5). Resulting bound fraction naturally assumes a hyperbolic tangent–like profile, with its midpoint determined primarily by the effective dissociation constant. In RBA or MM approximation the concentration of Ca^2+^-bound buffer can be presented as

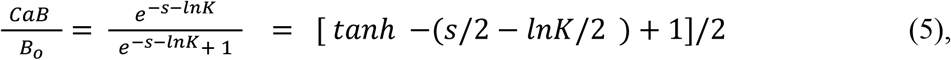

Thus the logarithm of dissociation constant serves as a mid-point of the wave of bound Ca^2+^. When K increases, its logarithm becomes more negative that leads to denser spacing of profiles, compare e. g. Fig. 2C vs. Fig. 2D.

### 3.2. One-dimensional propagation of Ca^2+^-bound species generated by single α-synuclein channels

To test the theoretical predictions experimentally, one-dimensional Ca^2+^ profiles were generated using outward currents through individual α-synuclein (α-Syn) channels incorporated into excised inside-out membrane patches. The channels are about equally permeable to K^+^, Na^+^, and Ca^2+^ (Mironov, 2015), but as the bath contains excessive Ca^2+^, the ‘outward’ recorded represents almost a pure Ca^2+^ current through single α-Syn channel. This preparation offers the advantage that channel openings frequently persist for several milliseconds, permitting direct observation of the temporal evolution of Ca^2+^-dependent fluorescence signals.

The patches were set at the potential that generated a pure Ca^2+^-current of 0.2 pA that match a typical single L-type channel current (Mironov & Richter, 1998) that correspond to [Ca^2+^]_o_/B_o_ = 0.1 for default set of parameters (Table 1). Opening of a single channel produced localized increases in Fluo-4 or Fluo-4-dextran fluorescence corresponding to the formation of Ca^2+^-bound indicator molecules. Kymographs revealed a progressive expansion of the fluorescence signal from the pipette tip toward the pipette interior. Line scans extracted at successive time points demonstrated that the fluorescence front propagated continuously away from the source rather than remaining spatially stationary.

The observed propagation is consistent with the predicted wave behavior of Ca^2+^-bound species. Although fluorescence decayed rapidly following channel closure because of diffusion and ongoing binding reactions, the spatial evolution during channel opening was well described by a propagating front. Mean estimates of experimental velocities are summarized in Table 2 and agree qualitatively with theoretical predictions.

### 3.3 Two-dimensional profiles of bound Ca^2+^ in neuronal soma

The theoretical predictions were further evaluated in a two-dimensional cellular geometry using cultured dorsal root ganglion neurons loaded intracellularly with Fluo-4 or Fluo-4-dextran. Whole-cell Ca^2+^ currents were evoked by depolarizing voltage steps, producing reproducible increases in intracellular fluorescence.

In the beginning of experiment, the voltage pulses produced some increase in the middle of the cell due to excitation of the basal membrane. When the cell was ‘trained’ with 100-long pulses delivered at 0.1 Hz to deplete interstitial space, this offset strongly diminished and the lateral transients that originate at cell borders were seen more clearly. Immediately after cell excitation, fluorescence first increased beneath the plasma membrane before progressively spreading into the bulk of the cytoplasm. Line-scan analysis demonstrates that the resulting Ca^2+^-bound fluorescence propagated inward as a traveling front.

Pharmacological manipulation of Ca^2+^ influx produced the changes predicted by the analytical model. Enhancement of L-type Ca^2+^ channel activity with Bay K 8644 increased current amplitude and accelerated propagation of the fluorescence front, whereas inhibition with Nifedipine reduced both current amplitude and wave velocity. These effects were observed for both indicator dyes but were particularly pronounced with the slowly diffusing dextran-conjugated probe. Front position changed proportional to √t, which is characteristic of diffusion process (a self-similarity variable is *z = x/2√t*, see Eqs. (2) above and (A3) below.

The experimentally determined propagation velocities (Table 2) followed the theoretical trends predicted by the WKBJ analysis: larger Ca^2+^ influxes produced faster-moving waves, whereas reduced influx slowed propagation. Moreover, the overall similarity between the one-dimensional and two-dimensional measurements supports the conclusion that the spread out of Ca^2+^-bound species represent a robust consequence of reaction–diffusion dynamics rather than a feature specific to a particular experimental geometry.

## 4 Discussion

The present study complements and extends two widely used concepts in biochemistry and biophysics: the Michaelis–Menten formalism (1913) and the Excess Buffer Approximation (EBA) introduced by Neher (1986). The Michaelis–Menten approach describes local substrate–enzyme interactions but does not account for the diffusion of either substrate or enzyme. Likewise, the EBA assumes that Ca^2+^ buffers are present in excess and therefore remain far from saturation. This assumption permits linearization of the original reaction–diffusion (RD) system (Eq. 1), reducing it to a second-order ordinary differential equation (Egs. (1) and (A11)). The resulting solution predicts the formation of localized Ca^2+^ nanodomains around persistently open Ca^2+^ channels. This concept is frequently invoked to explain a variety of physiological phenomena as recently reviewed by Eisner et al. (2023).

Most theoretical studies of intracellular Ca^2+^ signaling have focused on exponentially decaying steady-state Ca^2+^ distributions (Neher, 1986; Naraghi & Neher, 1997; Smith et al., 2001). However, these analyses critically depend on the assumption of excess buffer. When this constraint is removed, the analytical solution of irreversible Ca^2+^ binding for nearly equal Ca^2+^ and buffer diffusion coefficients, predicts appearance of periodic Ca^2+^ patterns under specific conditions (Mironov, 2019). Exact solutions were also found in the case of immobile buffer that bind Ca^2+^ irreversibly (Mironov, 2023a, no RBA in this case). An important conclusion emerging from these analyses is that free Ca^2+^ and Ca^2+^-bound species may exhibit markedly different dynamic behaviors. While free Ca^2+^ accumulates relatively slowly around an individual channel, Ca^2+^ bound to a buffer or sensor can propagate away from the source at substantially higher velocities (Mironov, 2023a). Consequently, the effective signal carriers—the Ca^2+^-bound buffer or sensor molecules—may travel both faster and farther than the signaling ion itself.

To fully understand the information conveyed by intracellular Ca^2+^ signals, it is necessary to consider not only the distribution of free Ca^2+^ but also the generation and propagation of Ca^2+^-bound buffers and sensors. Fluorescence measurements report Ca^2+^ bound to the indicator rather than free Ca^2+^ itself and the theoretical analysis therefore focused on the dynamics of the bound species. In the theoretical part of this study we used RBA that is identical to Michaelis–Menten kinetics extensively used in biochemistry. Wagner and Keizer (1994) were first to apply RBA to intracellular Ca^2+^ buffering and derived a nonlinear RD equation for total cellular Ca^2+^. Their analysis relies primarily on numerical solutions and limiting cases. However, the approximate analytical solutions are possible in RBA when logarithmic approximation of total is Ca^2+^ used (Mironova and Mironov, 2008).

The present study further extends and tackles this concept by explicitly incorporating diffusion into MM-type reaction kinetics. To derive analytical solutions, the Liouville–Green (LG) transformation was employed, as described in Sections 5.1 and 5.2. In quantum mechanics, this approach is widely known as the Wentzel–Kramers–Brillouin–Jeffreys (WKBJ) method and is commonly used to solve the stationary Schrödinger equation. Applications of WKBJ techniques to time-dependent reaction– diffusion systems are comparatively rare. Among the few examples are the studies of Krause et al. (2011), although these authors employed a stationary WKBJ formalism in which temporal dependence entered only indirectly.

To avoid overburdening the main text, all mathematical derivations and numerical analyses are presented in Sections 5.X. As an initial validation of the approach, the WKBJ formalism was applied to multidimensional diffusion and to diffusion coupled with a first-order reaction corresponding to the time-dependent version of the EBA (Section 5.1). The resulting solutions accurately reproduced established analytical results. Remarkably, the leading-order WKBJ approximation alone proved sufficient to achieve excellent agreement with exact solutions (Section 5.1; Fig. A).

The same methodology is subsequently extended to the Ca^2+^–buffer system (Section 5.2). The resulting derivations yield simple and transparent analytical expressions as summarized in Eqs. (2–4). Once again, the leading-order WKBJ term appeared sufficient to reproduce the behavior of multidimensional RD systems (Fig. 1). Although only a single buffer species was considered here, the formalism can readily be extended to multiple buffers with different mobilities, in a way previously proposed for related RBA models (Mironova & Mironov, 2008).

The theoretical analysis identifies buffer mobility as a major determinant of intracellular Ca^2+^ patterns (Fig. 1). Contrary to the predictions of the classical EBA, these patterns are not stationary, even under conditions of constant Ca^2+^ influx. Owing to progressive buffer saturation, free Ca^2+^ distributions continuously expand away from the channel source in both linear (Fig. 1A,B) and circular (Fig. 1C,D) geometries. Even more striking behavior is observed for Ca^2+^-bound species, which distributions spread out over substantially larger distances.

For buffer diffusion coefficients approaching that of free Ca^2+^ (Fig. 1B), the distributions of both free and bound Ca^2+^ remain spatially confined, thereby supporting the classical concept of Ca^2+^ nanodomains (Neher, 1986). This paradigm was used to interpret the measurements performed using synthetic fluorescent indicator dyes whose diffusion coefficients are approximately close to that of free Ca^2+^. Endogenous cytoplasmic Ca^2+^ buffers such as calmodulin are larger molecules and possess substantially lower diffusion coefficients; consequently, they are often treated as effectively immobile buffers (Zhou & Neher, 1993; Schwaller, 2010). Both the theoretical predictions presented here (Figs. 1 and 2) and the corresponding experimental observations suggest that, under conditions of low buffer mobility, Ca^2+^-bound buffer and sensor molecules are not confined to localized domains but instead propagate progressively throughout the cytoplasm.

## 5 A novel mathematical treatment of reaction-diffusion (RD) systems using WKBJ formalism

### 5.1 Time-dependent solutions of simple diffusion equations

Before applying the Liouville–Green/WKBJ formalism to Ca^2+^ buffering, we first validate the approach using diffusion and reaction–diffusion equations for which exact analytical solutions are available. This comparison demonstrates that the leading-order WKBJ approximation accurately reproduces the known analytical solutions and provides the foundation for the subsequent analysis. Consider a non-stationary (time-dependent) case for a simple diffusion

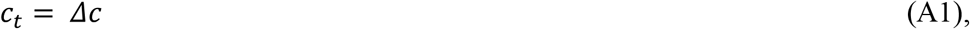

where subscripts *t* and *r* denote the time- and spatial derivative, respectively, and *Δ* represents the *n*-dimensional Laplacian

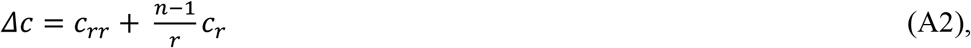

used in Eq. (1) (Crank, 1979; Carslaw & Jaeger, 1986). A straightforward way to obtaining the solution is to introduce the Boltzmann (Crank, 1979) or self-similarity variable *z = r/2√t* (Polyanin & Zaitsev, 2004). Eq. (A1) then transforms into the ordinary differential equation (ODE) as

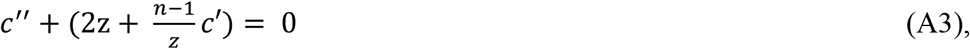

where primes indicate a derivative with respect to z. For further analysis, transform the equation into a canonical (Liouville-Green) form through the change of variable (Polyanin and Zaitsev, 2004)

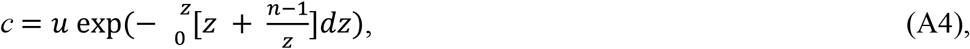

that gives

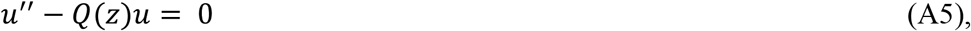

where

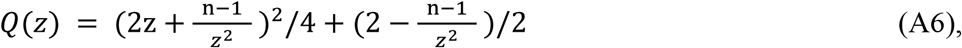

Eq (A5) is nothing else as a stationary Schrödinger equation with effective potential *Q*(*z*). Following Bender and Orzag (1999), we rewrite it as

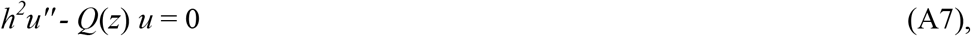

where *h* is a small parameter. A dependent variable is then presented as an exponential, whose argument is expanded in powers of *h* as

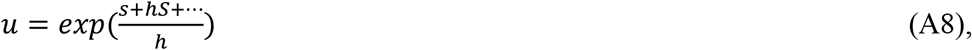

Next, substitute (A8) into (A7), divide off the exponential factors and collect the terms multiplied by the same power of *h*. The first two equations has solutions (Bender & Orzag, 1999)

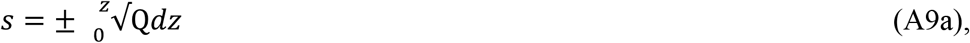

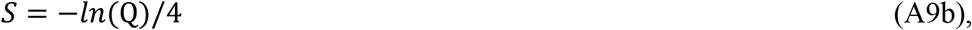

where s and S are the first and the second terms in WKBJ expansions (A8). In most WKBJ applications further correcting terms are small, and only two terms of expansion of exponent are fully sufficient for most purposes (Bender and Orzag, 1999). This is also valid in this study as shown below. For example, when Q(z) = *k*, the solution given by Eq. (A9a),

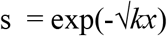

is identical to the analytical solution of ODE *c’’ = kc*, which appears as a steady state solution in EBA approximation (Neher, 1986).

For physically relevant solution only exponential with negative argument (-sign) in (A9a) is used. For multi-dimensional diffusion, the equations, potentials Q(z), and leading solutions *lnc* = s(*z*) are given in Table A. In 1D-case the integral (A9) is

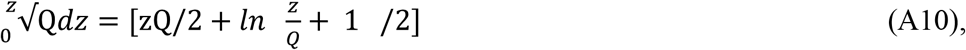

For circular diffusion, the direct integration of polynom *Q*(*z*) =√(z^2^ + 2 - 1/4z^2^) is possible, but a result is quite cumbersome, involving the elliptic integrals. Therefore an original equation (A7) was instead numerically integrated by the method of lines (Berezin and Zhidkov, 1965; Mironov, 2023a), briefly described in Section 5.3. In 3D case, the solution is simply 1D-result divided by *r*.

Fig. A shows that the leading WKBJ term (A9a) neatly reproduces exact solutions (left panels A, C and E), and the first order contribution (9b), did not modify the results substantially.

Next, consider the simplest RD equation

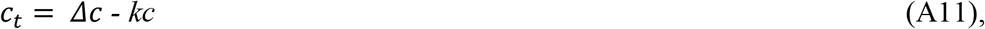

In comparison with pure diffusion it contains a spatially uniform first order reaction, whose rate constant is *k*. Eq. (A11) corresponds to the time-dependent excessive buffer approximation (EBA), which describes irreversible Ca^2+^ binding by buffer with rate constant *k = k*_*on*_*B*_*o*_ (Table 1). After transformation into to a self-similar form, the equation corresponds to (A3) with *4ktc* in the right-hand side. The solutions are readily obtained from (A10) in Table A with this term added to Q(z) determined for pure diffusion. The exact solution (Mironov, 2019) is

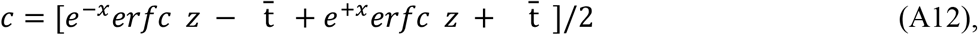

is again reproduced well by the leading term (Fig. A, right panels B, D and F).

### 5.2 Time-dependent analytical solutions of Ca^2+^ buffering using WKBJ formalism

In all derivations below we use the normalized equation derived from Eqs. (1) in Methods. There the relative buffer diffusion coefficient is *d = D*_*buffer*_*/D*_*Ca*_, and the intrinsic time and distance units are *τ = 1/k*_*on*_*B*_*o*_ = 50 μs and *r*_*o*_ *= √D*_*Ca*_*/B*_*o*_*k*_*on*_ = 105 μm, respectively (Table 1, see also Mironov, 2019, 2023a). The concentrations below are divided by the *total* buffer concentration *B*_*o*_ = 0.2 mM (Table 1). The system of RD equations for Ca^2+^ buffering is

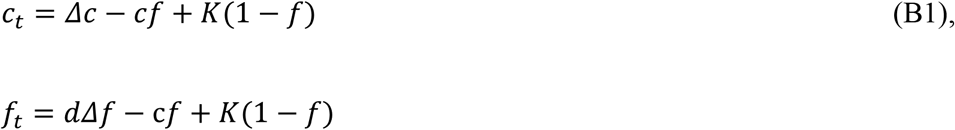

where *c, f* and *b* and define the concentrations of free Ca^2+^ and buffer, and bound buffer, respectively (all normalized to *B*_*o*_). Note that *b = 1 – f* . The normalized dissociation constant is *K = k*_*off/*_*k*_*on*_*B*_*o*_ = 0.003 (Table 1).

The first two terms describe the diffusion and the last two terms represent forward and backward steps in reversible reaction. It is convenient to reorganize a reaction term as

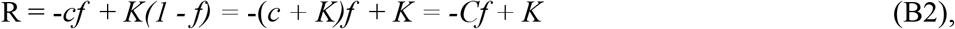

where a new Ca^2+^ dependent variable *C = c +K* is introduced.

In ‘multiple scales’ presentation (Bender & Orsag, 1999, see also Eq. (A7) above), we need to solve a system

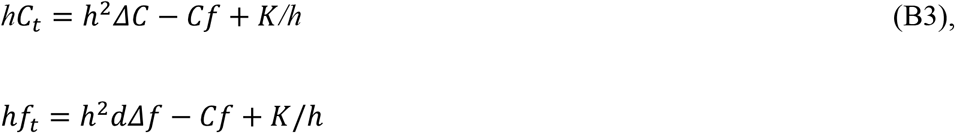

When the reaction term

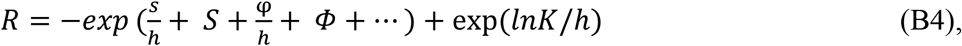

is expanded in exponentials in the leading order *h*^*-1*^, the reaction term *R = 0* and due to identity

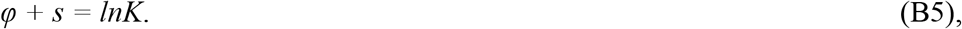

or

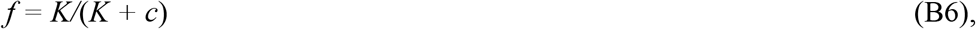

that describes the equilibrium between Ca^2+^ and buffer as presumed in RBA or MM formalism.

Expanding the concentrations as exponentials, *C = exp*(s*/h + S +*…) and *f = exp*(*φ/ h + Φ +*…) the system (B3) is written as

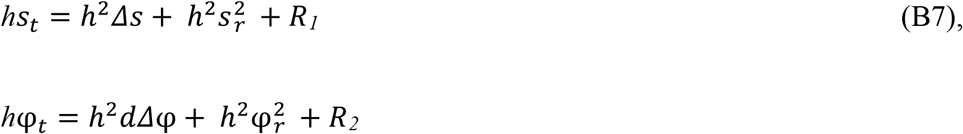

where the reaction terms are

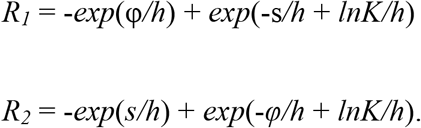

From (B5) follows that *R*_*1*_ *= R*_*2*_ = 0. Subtracting the two equations in (B7), we obtain a single equation

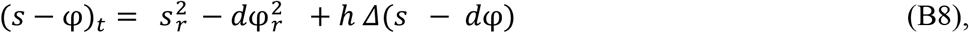

From (B5) also follows that for both time and space derivatives we have *φ’ = -s’* that simplifies (B8) to

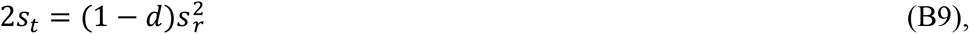

This is nothing else as a pure diffusion equation with apparent diffusion coefficient *D =* (*1-d*)/2. Its solution for different dimensions can be extracted from the Table A, line (A10), with *z* variable by scaled by √[(*1 - d*)/2]. Note that the case *d = 1* is critical (see also the end of Section 5.3 below) and should be treated separately using e. g. full Eq. (B1), whose solution was obtained previously (Mironov, 2019).

Subtraction of the two Eqs. (B4), in the next order *h*^*0*^ gives

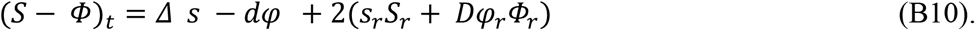

To express *Φ* through *S*, we use (B4) and (B6) to obtain

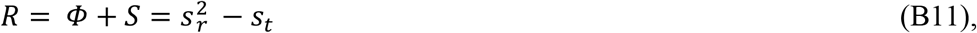

from which *Φ* derivatives can be defined as

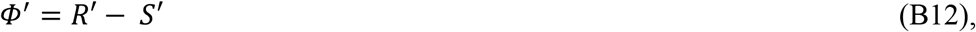

Inserting (B12) into (B10), we obtain the first order PDE that may be solved analytically (Polyanin et. al. 2002), but the explicit solution is rather cumbersome. Therefore (B12) was solved numerically. The data plotted in Fig. 1 in the main text indicate no significant modification of the solution given by the leading WKBJ term (B9).

Explicit formulas for leading WKBJ term in 1D-case formalism are listed in Section 2.6 in Methods, Eqs. (2-4). 3D-results can be readily obtained from 1D-case by dividing the solution by *r*. In the next Section and Fig. B is shown that WKBJ ansatz reproduces well numerical solutions of the full RD system (B1), indicating that WKBJ approach offers simple possibility to incorporate into MM (or RBA) formalism the effects of diffusion.

### 5.3 Application of the method of lines to solving RD equations

The algorithm is illustrated for a typical RD equation

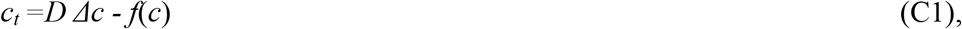

for diffusion in the presence of sink, whose concentration dependence may be non-linear. For pure diffusion the algorithm has been described by Berezin and Zhidkov (1965). Below it is presented following (Mironov, 2023b) with appropriate modifications for non-linear sink. Briefly, write down (C1) in discrete form as

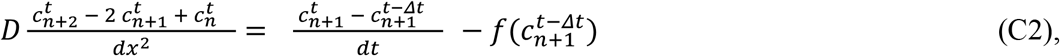

The solution proceeds in two steps. First we assume a linear relationship between 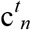 and 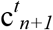

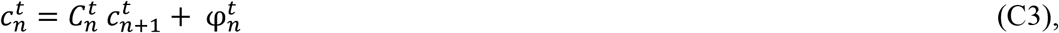

Setting *γ = D dt/dx*^*2*^, we present (C2) as

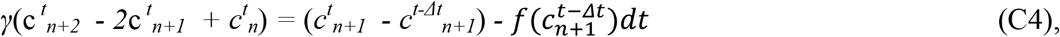

As in (C3), we can derive a linear connection between 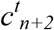 and 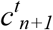 as

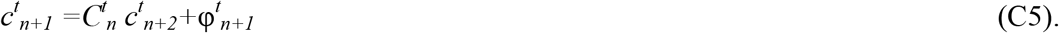

where

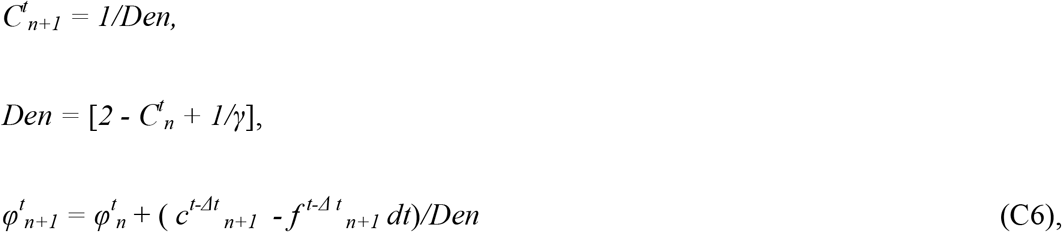

In numerical calculations one starts from initial conditions e. g.

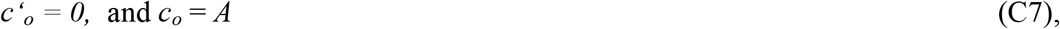

and subsequently evaluates all coefficients *C*_*n*_ and *φ*^*t*^_*n*_ via Eqs. (C6). Then we set a limiting value to *c*_*end*_ = 0 or another constant, and move backward to obtain all concentrations c_*n*_ at given *t* from Eq. (C3). The two progressions Eq. (C5) and (C6) explicitly define both the forward and backward steps that explains a name ‘chasing’ for the algorithm (Berezin & Zhidkov, 1965). The algorithm has been tested for many ODEs and accurately recovered exact solutions (Mironov, 2023b). In comparison with other approaches designed to solve RD equations the main advantage of the method is the possibility to work with bigger time steps without loosing accuracy and stability.

The method of lines was used to solve the full equation (B1). Fig. B presents the results of calculation for sample cases presented in Figs. 1 and 2 above. The solutions slightly disagree at *d* > 0.9, close to the critical value *d* = 1. In this case it is better to solve the full system (Mironov, 2019). Nonetheless, a nice correspondence between the results obtained analytically and numerically indicates that LG-WKBJ ansatz may be used as a simple analytical alternative to numerical solution of RD equations.

Theoretical velocities from front position at 12 ms from data shown in Figs. 1, 2. In the second column the numbers are obtained for basic (default) model parameters. In next columns the velocities are obtained in calculation runs, when of only one parameter in the basic set was modified as indicated in the first line. Experimental velocity estimates give the mean wave speeds ± SD as measured for Ca^2+^-bound buffer (organic indicator dye, fluo-4(X)) in 1D-(single α-Syn channel, Fig. 3) and 2D-systems (DRG neurons, Fig. 4). Experimental estimates were obtained from FWHM (midpoints) of linescans in Figs. 3, 4 divided by elapsed time. The data were analyzed for four different patches with α-Syn channels and four whole-cell patched cells. Control values vs. BAYK data had *P* values equal to 0.0006 (Fluo-4) and 0.0088 (Fluo-4-dextran) respectively.

**Fig. 3.**
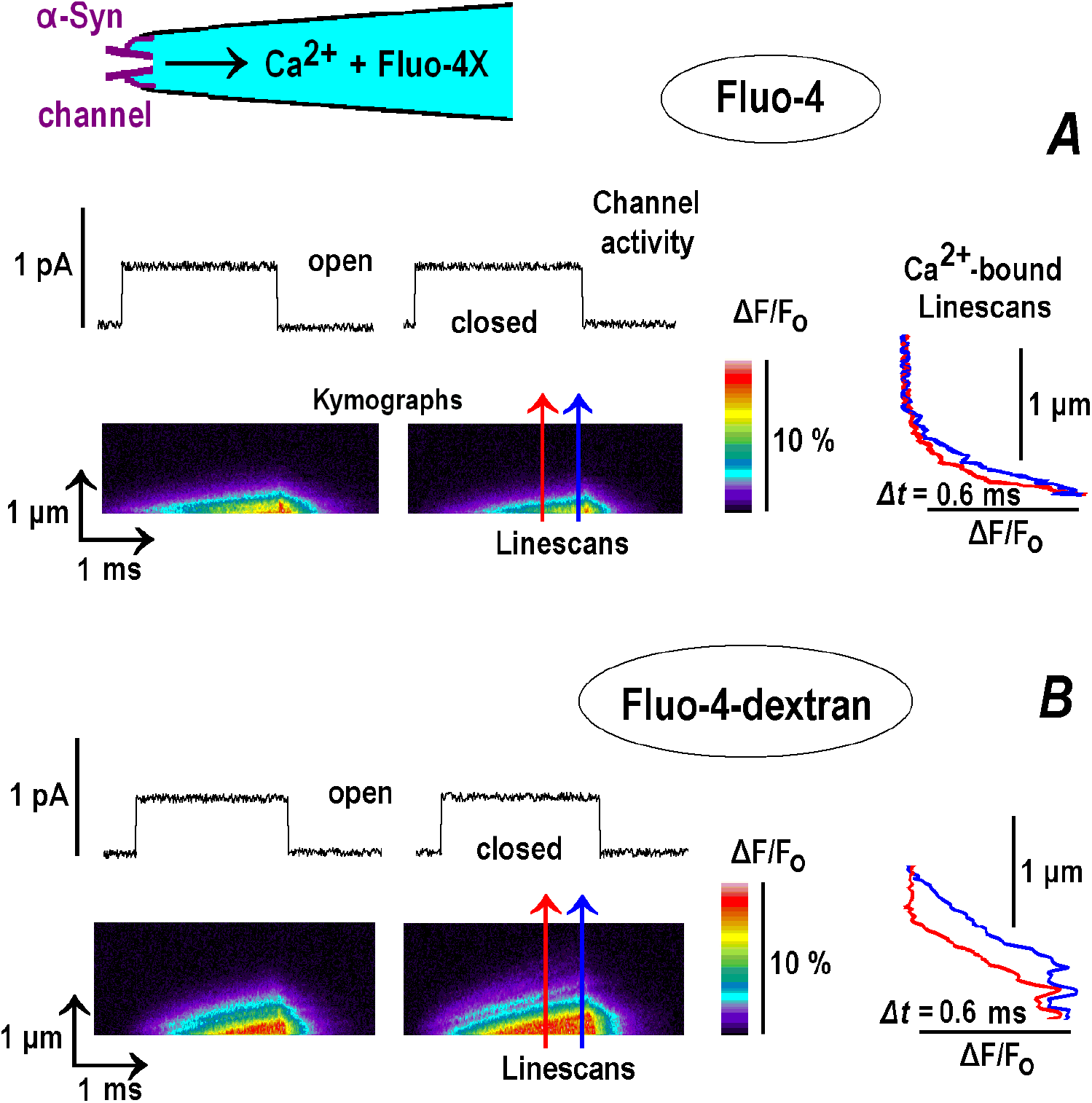
1D-Ca^2+^transients generated by α-synuclein (α-Syn) channel. Single channels were incorporated into inside-out patches excised from cultured hippocampal neurons. The pipette solution contained either fluo-4 (***A***) or fluo-4-dextran (***B***). The experimental configuration is sketched in the inset (top left). Channel opening induced ‘outward’ Ca^2+^ current (the bath contained 88 mM CaCl_2_) and related local increases of Ca^2+^ bound to the indicator dye in the pipette tip. The panels show sample events obtained in four patches for each indicator dye. Note that after channel closure, the fluorescence decay fast due to diffusion of Ca^2+^ and reaction with free buffer molecules in the bulk. The panels ***A*** and ***B*** show a single channel current (top) and the fluorescence of bound Ca^2+^ as kymographs (bottom) The relative changes in fluorescence are obtained from background subtracted images and plotted according to the color calibration bar in the middle. The time runs from left to right and the distance from the pipette tips runs upward. Increasing normalized fluorescence (*F/F*_*o*_) corresponds to an increase in bound Ca^2+^ (warmer colors). A monotonous increase in fluorescence indicates an expansion of bound Ca^2+^. Rightmost in both panels shown are the distance scans made as indicated by arrows of same color showing also the direction of scanning.

**Fig. 4.**
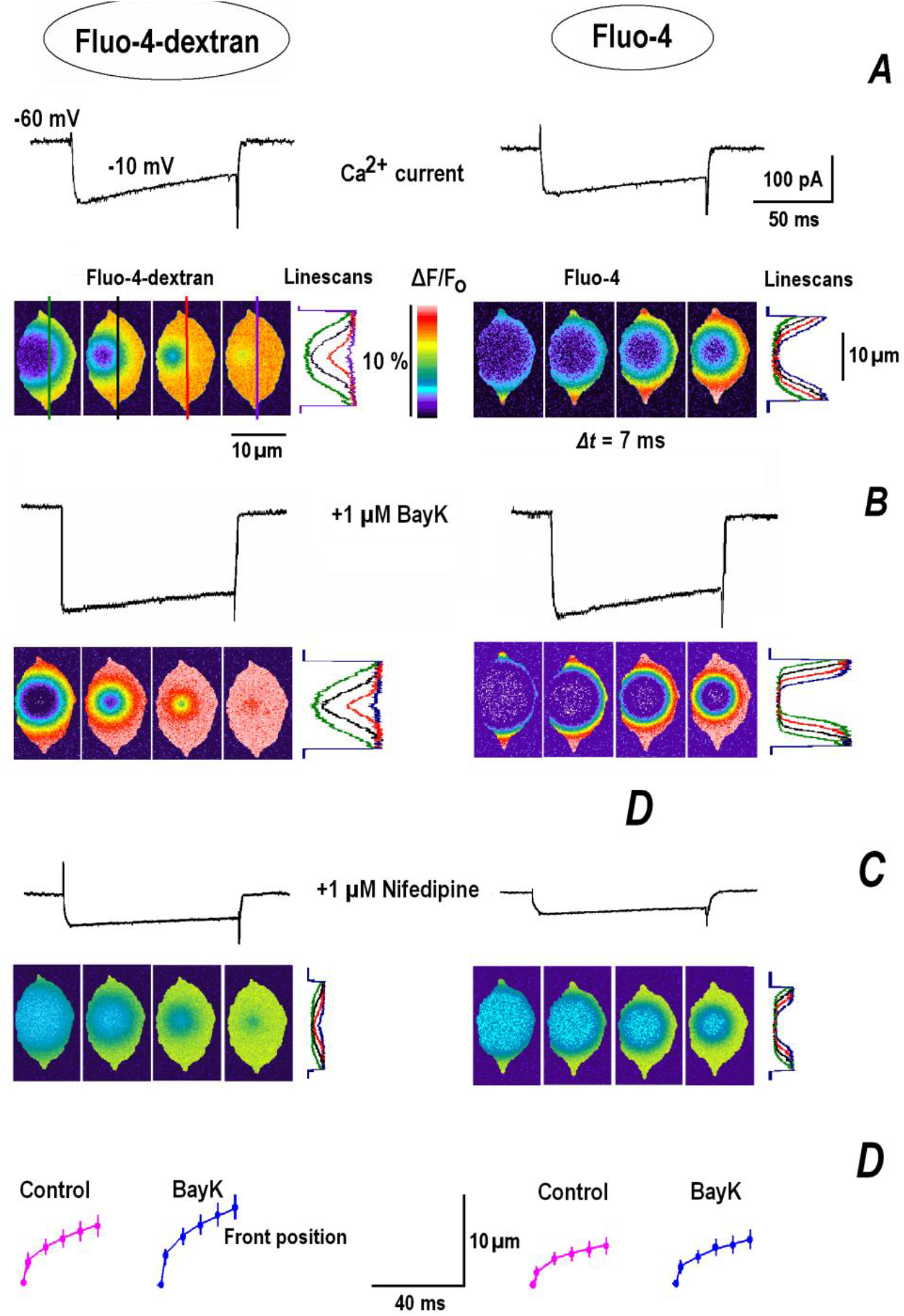
2D-’circular’ Ca^2+^ diffusion in dorsal root ganglion (DRG) neurons. Each panel shows whole-cell Ca^2+^-channel current evoked by step depolarizations by 100 ms-long pulse from -60 to -10 mV (top traces). The cell images (taken at interval 7 ms starting from 1.5 ms) were used to obtain the vertical linescans through the middle of images indicated as vertical lines with the same color in (***A***). The panels show the results obtained in the control (***A***), 5 min after addition of agonist of L-type Ca^2+^-channels (***B***) and its wash-out with antagonist (***C***). The last panel ***D*** shows the plots of measured front positions vs. time as means ± SD obtained from four cells for each indicator dye. The lines depict appropriately scaled √t-dependence indicative a diffusion process.

**Table A.**
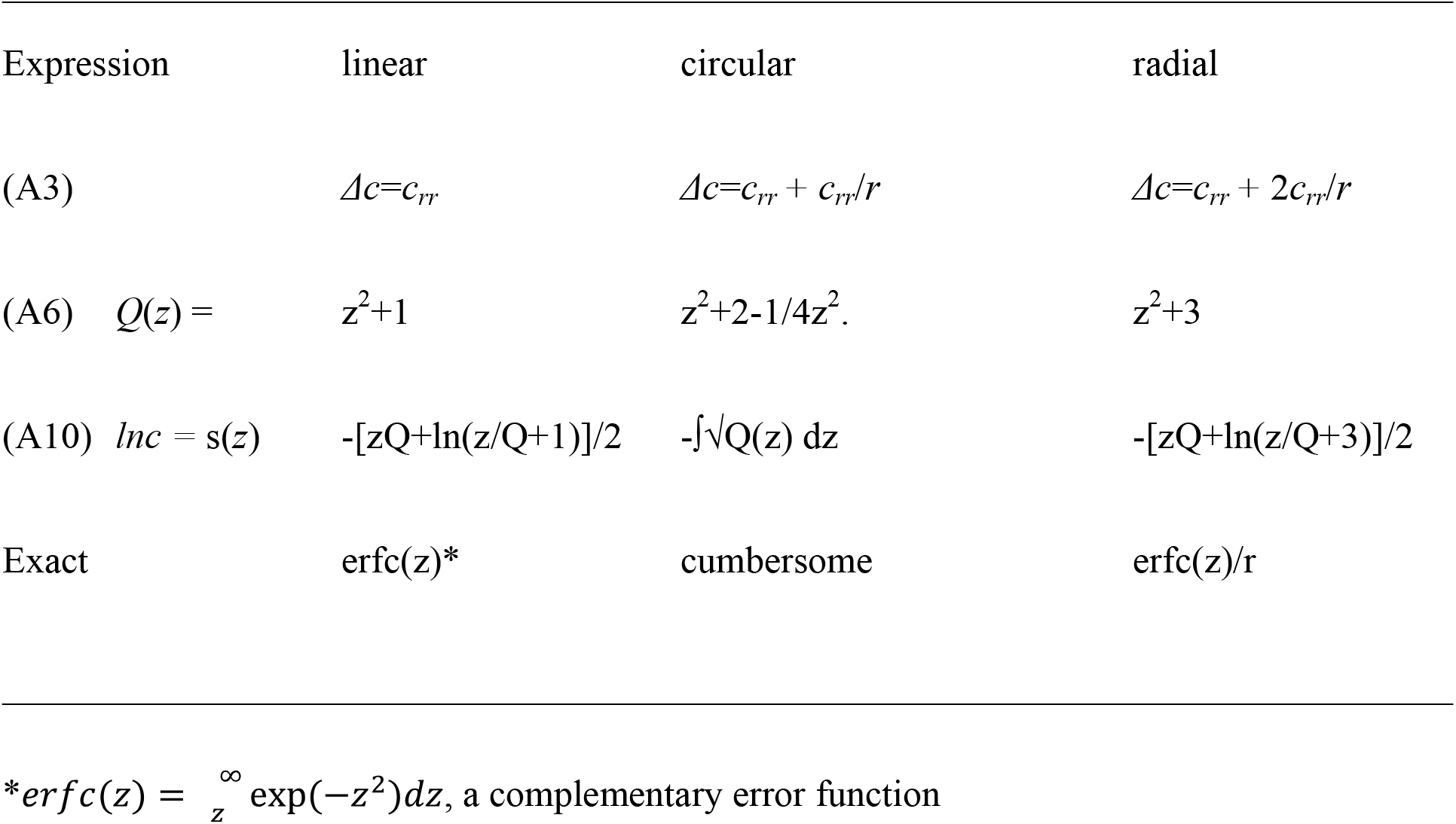
Explicit expressions of functions used in derivation of leading term in WKBJ solutions for multi-dimensional diffusion.

## Figures

**Fig. A.**
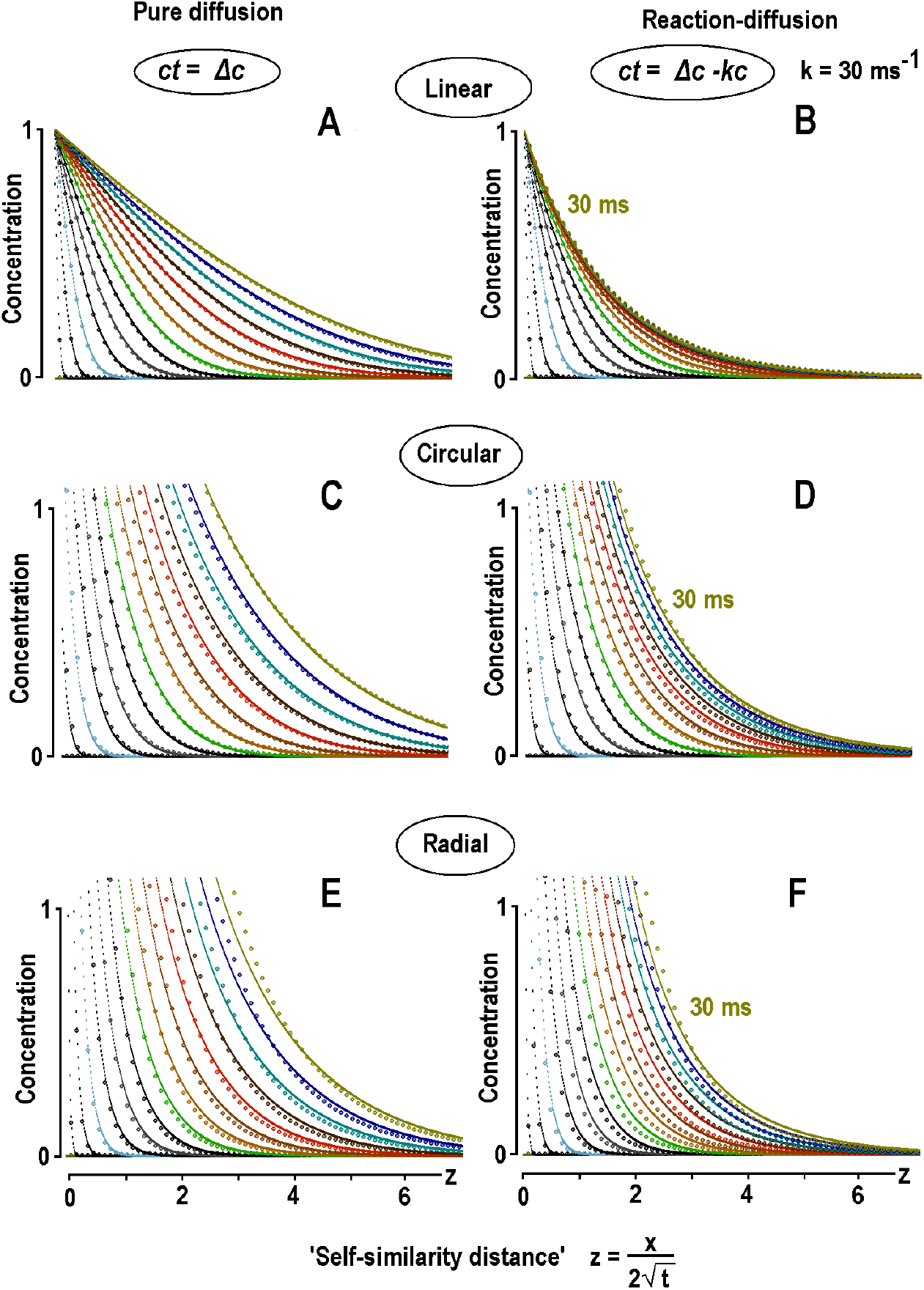
LG-WKBJ approach to solve reaction-diffusion PDEs. Solutions are obtained for multi-dimensional (1-, 2- and 3-D) diffusion (left panels) and reaction-diffusion equations (right panels) as indicated. Exact solution of diffusion (*c*_*t*_ *= Δc*) and reaction-diffusion equations (*c*_*t*_ *= Δc-kc, k* = 30 ms^-1^) are plotted as lines. Time-dependent changes given by established analytical solutions (lines) are well reproduced by the leading term in LG-WKBJ (dots). For linear and radial geometries the curves are calculated from explicit expressions listed in Table A, and for circular diffusion they are obtained by numerical integration of Eq. (A5). Concentrations (*y*-axes) are plotted vs. the Boltzmann variable (a self-similarity ‘distance’). In 30 ms-long runs, the time interval between the curves increased quadratically to obtain about equally separated curves. The profiles become sharper with increasing dimensionality, and the curves obtained in 2D- and 3D geometries are cut from above. Introduction of ‘diffusion sink’, which describes the bulk uptake with the rate constant *k*, decreases the width of profiles that ‘saturate’ with time (right row of panels).

**Fig. B.**
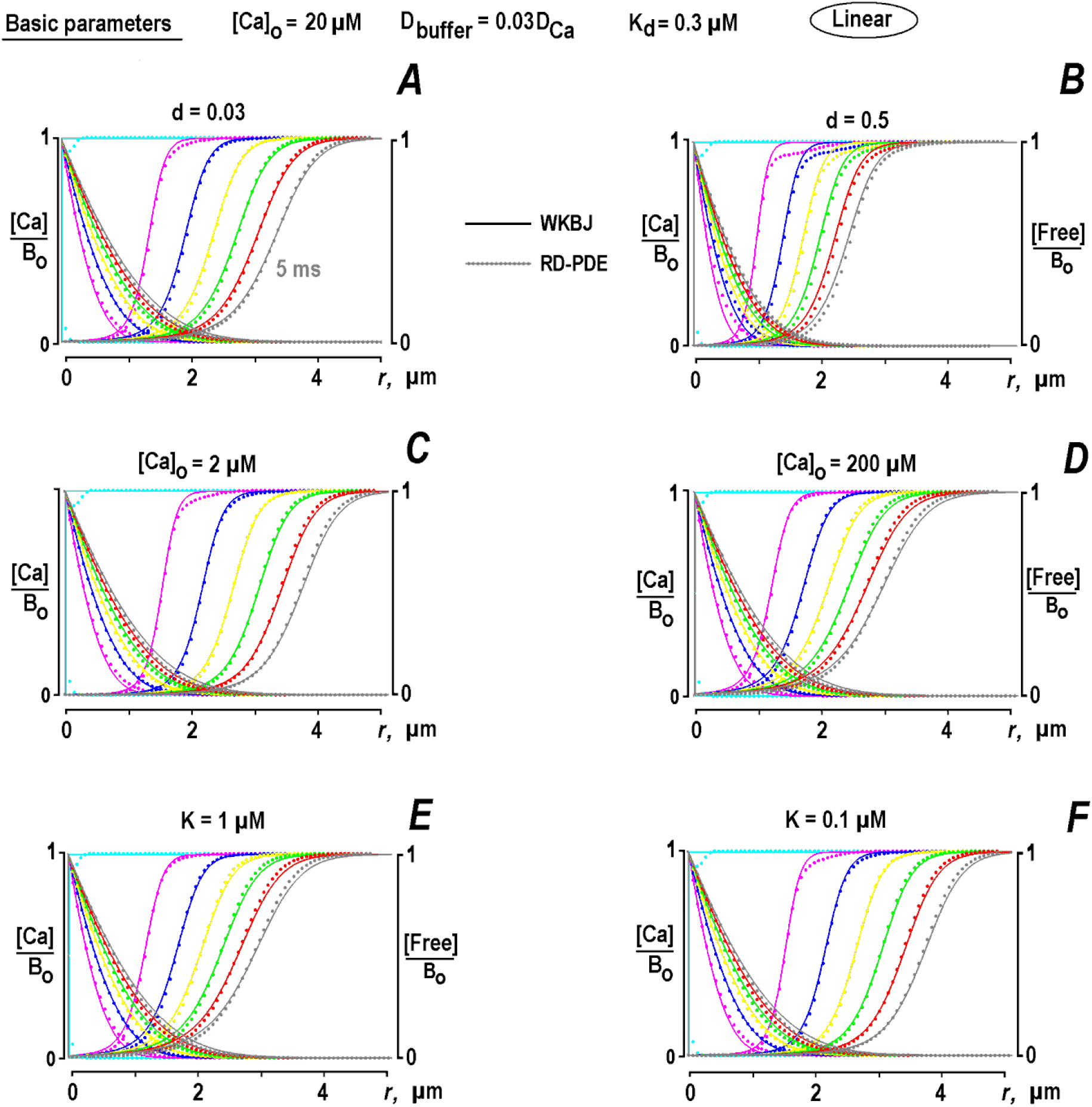
LG-WKBJ ansatz closely reproduces numerical solution of reaction-diffusion PDE. The full RD equation for Ca^2+^ buffering (B1) was solved by the method of lines (Section 5.3). The lines present the leading term in LG-WKBJ ansatz (lines), and the dots depict numerical solutions of full system (B1). In ***A*** the calculations were made with ‘default’ parameters (Table 1) and in other panels only one key parameter was changed as in Fig. 2. Only linear case was examined and the results in 2D-geometry show similar trend. The data indicate that LG-WKBJ approach provides a good analytical alternative to numerical approach in solving RD equations.

## Funding

The work was supported by DFG grant MI 685/2-1 and funded by Deutsche Forschungsgemeinschaft through the DFG-Research Center for Molecular Physiology of the Brain.

## Conflict of Interest

The author declares no competing interests.

## Author contributions

S.L.M. developed the concepts, mathematics, made all calculations, experiments, analysis, drafted and edited the manuscript.

## Notes

### Competing Interest Statement

The authors have declared no competing interest.

### Summary of Updates

After initial reviewing and submissio. No basic changes made

